# Age-Related Differences in Bimanual Coordination Are Associated with Increased Cerebellar Activity and Reduced Frontal Recruitment

**DOI:** 10.64898/2026.07.07.736910

**Authors:** Anastasia Sofia Weakley, Mikael Novén, Kristoffer Hougaard Madsen, Jesper Lundbye-Jensen, Hartwig Roman Siebner, Anke Karabanov

**Affiliations:** Human Movement Sciences, Department of Nutrition, Exercise and Sports, University of Copenhagen, Denmark; Department of Clinical Medicine, Interacting Minds Centre (IMC), Aarhus University, Denmark; Danish Research Centre for Magnetic Resonance, Department of Radiology and Nuclear Medicine, Copenhagen University Hospital - Amager and Hvidovre, Copenhagen, Denmark; Department of Clinical Sciences Lund, Division of Logopedics, Phoniatrics and Audiology, Lund University, Sweden; Department of Applied Mathematics and Computer Science, Technical University of Denmark, Denmark; Department of Radiology, Zealand University Hospital, Denmark; Department of Neurology, Copenhagen University Hospital Bispebjerg and Frederiksberg, Copenhagen, Denmark; Department of Clinical Medicine, Faculty of Health and Medical Sciences, University of Copenhagen, Copenhagen Denmark

## Abstract

Bimanual coordination declines in late adulthood, but the neural mechanisms underlying these changes remain unclear. Age-related differences in brain activity have been interpreted either as compensatory recruitment of frontal cognitive control regions or as a shift toward feedback-based control, supported by sensory and cerebellar processing systems. To investigate these hypotheses, we examined brain activity, using fMRI in twenty-three younger and twenty-three older adults performing a bimanual visuomotor pinch-force task with different task complexities.

Behaviourally, older adults showed lower accuracy than younger adults, particularly when task demands increased. Neuroimaging results revealed general age-dependent increases in activity within posterior cerebellar lobules VI–VII, regions overlapping with the classical oculomotor vermis and implicated in visuomotor adaptation, movement calibration, and error-based motor learning. In addition, during the more demanding task condition, older adults showed a greater increase in activation of anterior cerebellar lobules IV–V and a decrease in activation of the medial frontal pole (BA10). No consistent age-related increases or decreases in task related activation was observed in parieto-frontal regions. Moreover, better task performance across age groups was associated with lower activation in frontal cognitive control regions, including the superior medial frontal gyrus and right inferior frontal gyrus.

Together, these results suggest increased feedback- and error-related sensorimotor processing in older adults involving the cerebellum and frontal cortex.

## 1. Introduction

Many daily activities require coordinated bimanual motor control, but this ability deteriorates in late adulthood. Older adults typically perform bimanual movements more slowly and with reduced accuracy than younger adults do (Fling & Seidler, 2012; Roman-Liu & Tokarski, 2021). These age-related differences increase with higher task complexity, and motor performance of bimanual tasks, that require asynchronous coordination between the hands, is particularly vulnerable to decline (Kang et al., 2022; Maes et al., 2017; Zvornik et al., 2024).

At the neural level, functional magnetic resonance imaging (fMRI) studies have shown a mixed pattern of hyper- and hypoactivation during bimanual performance in older adults compared to younger adults (Zapparoli et al., 2022). Increased activation in older adults is commonly reported in the supplementary motor area (SMA), cerebellum, parietal and occipital cortex, and medial and dorsolateral prefrontal regions, whereas reduced activation is observed in the sensorimotor cortex surrounding the pre- and postcentral gyri and in subcortical structures, including the cerebellum, thalamus, and basal ganglia (Coxon et al., 2010; Goble et al., 2010; Monteiro et al., 2017). The cerebellum, in particular, shows both hyper- and hypoactivation across studies, a pattern that is likely driven by differences in task demands. Increased cerebellar activation has been interpreted as reflecting greater reliance on motor learning and error-correction processes when movements become less automatic with age, whereas reduced activation may occur in studies reporting a shift from subcortical to cortical recruitment (Monteiro et al., 2017; Zapparoli et al., 2022).

Several influential neurocognitive aging models have been proposed to explain age-related changes in brain activation; however, none adequately account for the patterns observed in bimanual motor control. The Hemispheric Asymmetry Reduction in Older Adults (HAROLD) model was developed primarily for cognitive tasks and unimanual movements and does not readily generalize to bimanual coordination, where older adults can show increased rather than reduced lateralization (Cabeza, 2002; Wulff-Abramsson et al., 2025). The Posterior–Anterior Shift in Aging (PASA) hypothesis predicts frontal compensation for reduced posterior activity; however, bimanual control studies frequently report hyperactivation in the parietal and occipital cortices, contradicting a general poste-rior–anterior shift (Zapparoli et al., 2022). The Compensation-Related Utilization of Neural Circuits (CRUNCH) hypothesis predicts an altered activation–demand relationship in aging, with age-dependent over-recruitment during simple tasks, followed by early plateauing of neural signalling at higher task demands in older adults (Goble et al., 2010; Reuter-Lorenz & Cappell, 2008). However, evidence for CRUNCH in bimanual motor control is also weak, as several studies have shown that both older and younger adults upregulate the BOLD signal proportionally with increasing complexity (Goble et al., 2010; Van Ruitenbeek, 2023).

An alternative explanation for the observed age-related differences in brain activity is that they reflect shifts in motor control strategy rather than compensatory over-recruitment per se. Age-related increases in sensory noise and reductions in neural efficiency may impair feedforward-based movement initiation, leading to greater reliance on feedback-mediated control supported by internal prediction (Newell et al., 2009). Electroencephalography studies are consistent with a shift away from motor planning toward greater reliance on online, error-driven corrections with aging (Yordanova et al., 2024; Wulff-Abramsson et al., 2025). However, the limited spatial resolution of EEG prevents clear dissociation of areas related to anticipatory control processes from feedback-related activity in regions such as the cerebellum (Seidler et al., 2004).

Importantly, most fMRI studies of bimanual motor control in aging have relied on cyclic tasks with a stable spatiotemporal structure and minimal demands for trial-by-trial replanning. Consequently, neural processes related to anticipatory planning may be underrepresented. Here, we address this gap using a bimanual visuomotor pinch-force tracking task that requires continuous integration of anticipatory planning and sensorimotor feedback (Karabanov, 2023). Performance in this task is age sensitive, with older adults showing greater decline than younger adults when shifting from simple symmetric to more complex asymmetric bimanual pinch force contractions (Zvornik et al., 2024). This study aims to investigate whether age-related activation differences primarily reflect a shift in motor control strategy, from feedforward-based movement initiation toward feedback-mediated motor control. If so, older adults should show increased activation of cerebellar and fronto-parietal regions that become more pronounced with higher task complexity, as performance errors increase, rather than the early activation plateau predicted by CRUNCH.

## 2. Methods

### 2.1. Participants

Twenty-five young (22-27 y.o.) adults and twenty-nine older (65-70 y.o.) adults participated in this study. All participants were screened for dementia using the Montreal Cognitive Assessment (MoCA) tool (Nasreddine et al., 2005). Eight participants ended up being excluded for the following reasons: incidental findings (2), MoCA score < 26 (3), claustrophobia (1), and faulty scan (2). Therefore, the study ended up with twenty-three participants in the young group and twenty-three participants in the older group (demographics are presented in Table 1). All participants were right-handed as measured by the Edinburgh Handedness Inventory (EHI) (Oldfield, 1971), had no contraindications for MRI-scanning, had no personal history of diagnosed neuropsychiatric diseases, and were not taking any neuroactive medication. All participants received oral and written information about the study, gave written informed consent before participation, and were reimbursed for their time. The study was approved by the Regional Committee on Health Research Ethics of the Capital Region in Denmark (De Videnskabsetiske Komitéer-Region H; Journal-nr: H-21025010). Participants were recruited via an announcement on forsøgsperson.dk.

**Table 1:** Descriptive characteristics of the included participants. Reported values are group averages ± standard deviation (SD). Tested outcome measures include: EHI = Edinburgh Handedness Inventory; pinch force = maximal pinch grip strength; pegboard assemblies = number of assemblies completed on the Purdue Pegboard test; 30-s Sit-to-Stand = lower-limb functional performance; MOCA = Montreal Cognitive Assessment; VO₂ max estimation = estimated maximal oxygen uptake, reflecting cardiorespiratory fitness. The groups significantly differed in age, VO_2_ max, MOCA score, and pegboard assemblies. Note that VO_2_ max values are within the normal ranges for each age group (20-29 y.o. Male: 54.4 ± 8.4, Female: 43.0 ± 7.7; 60-69 y.o. Male: 39.2 ± 6.7, Female: 31.1 ± 5.1). An asterisk (*) indicates significant group differences.

|  | Young group (22-27 y.o.) | Older group (65-70 y.o.) |
| --- | --- | --- |
| **Sex (F/M)** | 11/12 | 11/12 |
| **Age (years - range) \*** | 24.7 (22 – 27) | 67.3 (65 – 70) |
| **EHI (score)** | 100.0 ± 0.0 | 99.3 ± 3.2 |
| **Height (cm)** | 176.6 ± 11.3 | 171.4 ± 6.8 |
| **Weight (kg)** | 73.9 ± 14.7 | 74.7 ± 14.0 |
| **BMI (kg/m<sup>2</sup>)** | 23.6 ± 3.6 | 25.4 ± 4.4 |
| **Pinch force Left hand (N)** | 86.3 ± 23.5 | 78.5 ± 20.6 |
| **Pinch force Right hand (N)** | 90.2 ± 21.6 | 82.4 ± 25.5 |
| **Pegboard Assemblies (#) \*** | 30.0 ± 5.6 | 21.2 ± 5.9 |
| **30s Sit-To-Stand (# repetitions) \*** | 17.8 ± 3.6 | 17.2 ± 5.4 |
| **MOCA score** | 29.0 ± 0.9 | 28.1 ± 1.3 |
| **VO<sub>2</sub> max estimation (mL/kg/min) \*** | 46.9 ± 8.4 | 32.1 ± 7.1 |

### 2.2. Experimental procedure

Before signing the consent form, participants were given a general description of the study and informed that they could withdraw at any time. They were then asked to complete questionnaires and screening forms (including the MoCA and EHI). Pinch force (Baseline® Lite Hydraulic Pinch Gauge Prod. Nr: 12-0226), 30 second sit-to-stand (S2S), estimation of VO_2_ max level using Seis-mofit® (VentriJect 2022, 2900 Hellerup, Denmark), and BMI measurements were obtained outside the scanner. The participants were then asked to perform the Purdue pegboard test. Before being placed in the scanner, participants were given a description of the bimanual task that they would perform. After this, they were asked to lie down in a supine position on the extendable table of the scanner and were given force sensors, one for each hand. An MRI-compatible pulse oximeter and a pneumatic respiration belt were placed on their little finger and chest, respectively, to measure their heart rate and breathing. The head coil was placed over their face with a small mirror attached, through which they could view the screen. Before starting the bimanual task, maximal voluntary contraction (MVC) was measured using the force sensors, and the participants were instructed to pinch the force sensors, one hand at a time, as hard as they could. MVC measurements were repeated three times for each hand, and the average MVC for each hand was used to calculate the force levels during the experiment. The force range during the experiment varied between 2.5-10% of the participants’ individual average MVC. Once the MVC had been calculated, the experimental task was performed across 4 blocks of fMRI scans, with each block consisting of an initial baseline condition (static hold at 5% MVC for 20 s), followed by twelve alternating sets of symmetric and asymmetric trials, with each set containing 20 trials that lasted 2 s each (Figure 1B). Participants were allowed short breaks within blocks between every set of 20 trials (lasting 5 seconds) as well as between blocks. After completing the four blocks of task fMRI, anatomical and diffusion-weighted MRI scans were obtained. The total scanning time was approximately 72 mins.

**Figure 1.**
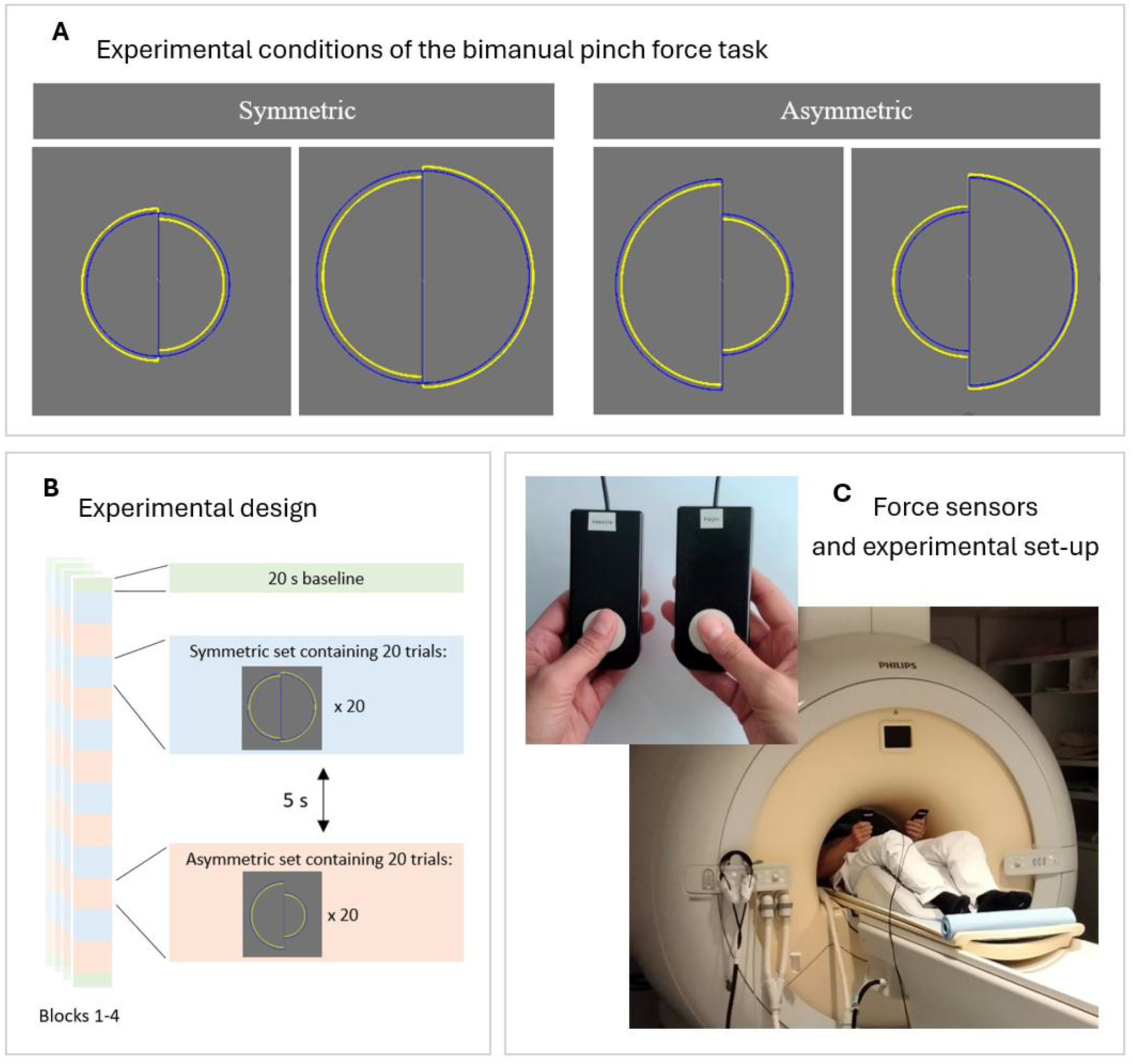
Illustration of the experimental design. **A:** Visually cued bimanual pinch force task. The aim was for the participants to align the yellow force-controlled semicircles with the blue target lines. The task involved two conditions: mirror symmetric, where both hands changed pinch force in the same direction (left), and inverse-asymmetric, where both hands changed pinch force in opposite directions (right). Target lines were variations of four different force levels: 2.5%, 5%, 7.5% or 10% of the average maximal voluntary contraction (MVC). **B:** The experimental procedure consisted of 4 blocks of fMRI, each containing 6 sets of symmetric and 6 sets of asymmetric trials, as well as baseline conditions. **C:** Participant with force sensors in the scanner.

### 2.3. Bimanual task

This study used a visually guided bimanual pinch force task. The participants held the force sensors between the thumb and index finger of each hand, allowing independent modulation of the pinch force. The force data were sampled at 1024 Hz, and the displayed force trace was smoothed by averaging the most recent 100 samples to reduce electrical noise.

The pinch force output from each hand was presented visually as a semicircle (dynamically updating at the screen refresh rate of 60Hz), with the right and left hands controlling the corresponding semicircles. Increasing pinch force increased the radius of the semi-circle, whereas decreasing pinch force reduced it. The task goal was to match the yellow force-controlled semicircles to the blue target lines displayed on the screen (Figure 1A). The participants were instructed to reach the target force as quickly as possible and then maintain it as accurately as possible for the remainder of each trial.

Each block included an initial baseline condition in which the participants produced a constant isometric pinch force for 20 s. This was followed by two dynamic bimanual conditions: a mirror-symmetric condition, in which both hands tracked the same force level and an inverse-asymmetric condition, in which the force levels differed between hands. The block concluded with a baseline condition (Figure 1B).

### 2.4. Image acquisition

Images were acquired at the Danish Research Centre of Magnetic Resonance (DRCMR), Hvidovre Hospital, Copenhagen, Denmark, using a 3T MR scanner (Philips Magnetom Achieva, Best, Netherlands) equipped with a 32-channel array receiver coil. Each scanning session included a T1-weighted image (MPRAGE; FOV: 245 mm; 245 sagittal slices, TR/TE: 6.0 / 2.7 ms; resolution 0.85×0.85×0.85 mm^3^; flip angle: 8 deg.; TI: 755.9 ms), four T2*-weighted echo planar imaging (EPI) sequences utilizing gradient echo (FOV: 190 mm; 42 transverse slices acquired in interleaved order, TR/TE: 1900/30; in-plane resolution: 3×3 mm2, 3mm slices, no slice gap, flip angle: 90 °, SENSE factor = 2), interleaved with four three volume EPI sequences with reversed phase encoding for susceptibility distortion correction and otherwise identical to the main EPI sequences, a T2-weighted image (TSE; 245 mm; 245 sagittal slices, TR/TE: 2.5 s / 270 ms; resolution 0.85×0.85×0.85mm^3^; flip angle: 90 deg.; TI: 755.9 ms), a Fluid-Attenuated Inversion Recovery image (FLAIR; 256 mm; 202 sagittal slices, TR/TE: 4.8 s / 330 ms; resolution 1×1×1mm^3^; flip angle: 90 deg.) and one diffusion-weighted sequence (FOV: 224 mm; 66 slices, 48 directions; TR/TE: 7970 /90; resolution: 2×2×2mm^3^, flip angle: 90 deg. b = 2000s/ mm2) with additional 5 b=0s/mm2 volumes acquired using reversed phase encoding for susceptibility error correction. FLAIR images were used for pathology assessments and for calculating white matter hyperintensity load. The analyses of the structural MRI data will be reported elsewhere.

### 2.5. Analysis of behavioural data

The time on target (ToT) was used as a measure of accuracy for the bimanual task. It was calculated as the percentage of trial time spent within 0.5% MVC of the force target lines for each trial. The resulting trial-level ToT values were entered into the statistical analysis, yielding multiple observations per participant. To test if there were any statistical differences in time on target (ToT), a linear mixed effects model was implemented in R (https://www.r-project.org/) using the R-package lme4 (Bates et al., 2015). Age, task condition, and hand were set as fixed effect variables, while the participant was set as a random effect variable. Significant main effects and interactions were estimated using the lmerTest R-package, which provides p-values from linear mixed-effects models using Satterthwaite’s degrees of freedom method (Kuznetsova et al., 2017).

### 2.6. Analysis of neuroimaging data

The data were pre-processed using fMRIPrep 22.1.1 (Esteban et al., 2018; Esteban et al., 2019), which is based on Nipype 1.8.5 (Gorgolewski et al., 2011; Gorgolewski et al., 2018). A full description of the pre-processing steps is provided in the Supplementary Material (S1). Briefly, the data were susceptibility distortion and motion corrected, slice timing corrected, and aligned to the MNI space prior to statistical analysis. Within-subject and between group statistical analyses were performed using SPM12 (Wellcome Trust Center for Neuroimaging, London, UK) using the mass-univariate general linear model (GLM). Prior to statistical analysis, the pre-processed data were smoothed using an 8 mm FWHM Gaussian kernel. The standard SPM canonical haemodynamic response function was used to generate a model for the design matrix, consisting of the bimanual context condition (symmetric and asymmetric) and the residual motion parameters from the rigid body realignment procedure in the pre-processing (6 parameters). In addition, the aliased harmonic expansion of pulse (6 regressors) and respiration (4 regressors) were added to model physiological noise (Glover et al., 2000). In order to control for age group differences in cerebrovascular health, white matter hyperintensity volume, calculated from the FLAIR image using the Sequence Adaptive Multimodal SEGmentation (SAMSEG) tool implemented in FreeSurfer (Puonti et al., 2016), normalized to total brain tissue volume, was also added to the model. Finally, the target levels of the left hand were added to decouple the response to changing bimanual context (asymmetric and symmetric) from the step-size of change between two target levels. Because these target levels were either perfectly correlated (symmetric condition) or perfectly anticorrelated (asymmetric condition), only the force level from one hand was required in the model. Contrast images from the first-level analyses were entered into random-effects group analyses to allow population-level inference. At the second level, a flexible factorial design was specified with the factors Age Group (Younger, Older) and Task Condition (Symmetric, Asymmetric). The primary effects of interest were the main effect of Age Group, the main effect of Task Condition, and their interaction. For all GLM analyses, multiple comparisons were corrected using SPM12’s family-wise error (FWE) correction method (p<0.05). The cluster defining threshold was set to p<0.001, uncorrected.

We investigated whether task-related activity covaried with behavioural performance in any brain regions. Here, the average ToT for each participant in the two task conditions was correlated with the beta values for the corresponding fMRI task condition contrast. We initially ran the analysis separately for the groups and task conditions, resulting in four GLMs. However, we found no significant correlation between brain activity and task performance in these models. Therefore, we also ran the analysis for averaged task conditions and age groups, using normalized ToT scores, which resulted in one combined GLM with three significant clusters. To account for age-group differences in overall task performance, ToT scores were standardized (z-scored) separately, within each age group, prior to pooling the data. This procedure removed between-group mean differences while retaining individual variability within each group.

For all GLM analyses, we used an exclusive ventricle mask, and significant clusters were labelled using an Automated Anatomical Labelling atlas (AAL) (https://www.gin.cnrs.fr/en/tools/aal/) implemented in SPM12 and confirmed manually with an MRI brain atlas (Cho et al., 2010). For labelling of cerebellar activation, we also used the JuBrain Anatomy Toolbox (Eickhoff et al., 2005), as it contains a detailed cerebellar atlas (Diedrichsen et al., 2009).

## 3. Results

### 3.1. Behavioural performance

Figure 2 shows the performance of the two groups during the symmetric and asymmetric task conditions. A higher score indicates better task performance (greater proportion of time spent on target). Younger participants performed the bimanual task more accurately in general than older participants (Main effect of Group: F(1,44) = 46.976; p<0.0001; β = 9.8; 95% CI = [0.067, 0.129]), and both groups performed the symmetric task condition more accurately than the asymmetric task condition (Main effect of Task: F(1,88259) = 4181.311; p<0.0001; β = 10.8; 95% CI = [0.102, 0.115]). Both groups also performed both tasks more accurately with their right hand compared to their left hand (Main effect of Hand: F(1,88259) = 487.884; p<0.0001; β = 2.7; 95% CI = [0.025, 0.03]). In addition, there was a significant interaction between task and age group (Interaction Group x Task: F(1, 88259) = 293.736; p<0.0001; β = −2.4; 95% CI = [-0.034, −0.014]), with higher performance drops during the asymmetric task in the old group. Post-hoc tests revealed a significant difference between old and young adults in the symmetric condition (p<0.0001) and asymmetric condition (p<0.0001), however the magnitude difference between the age groups was greater in the asymmetric condition (Table 2). No other significant interactions were observed.

**Table 2.** Mean ToT scores (% of trial time) for each age group and task condition. Standard deviation in brackets. Magnitude differences for each task condition showing the difference in mean ToT scores between age groups.

| ToT | Young | Old | Magnitude difference |
| --- | --- | --- | --- |
| Symmetric | 43.0<br>(18.7) | 34.6<br>(19.7) | 8.4 |
| Asymmetric | 37.2<br>(19.8) | 24.5<br>(19.6) | 12.7 |

**Figure 2.**
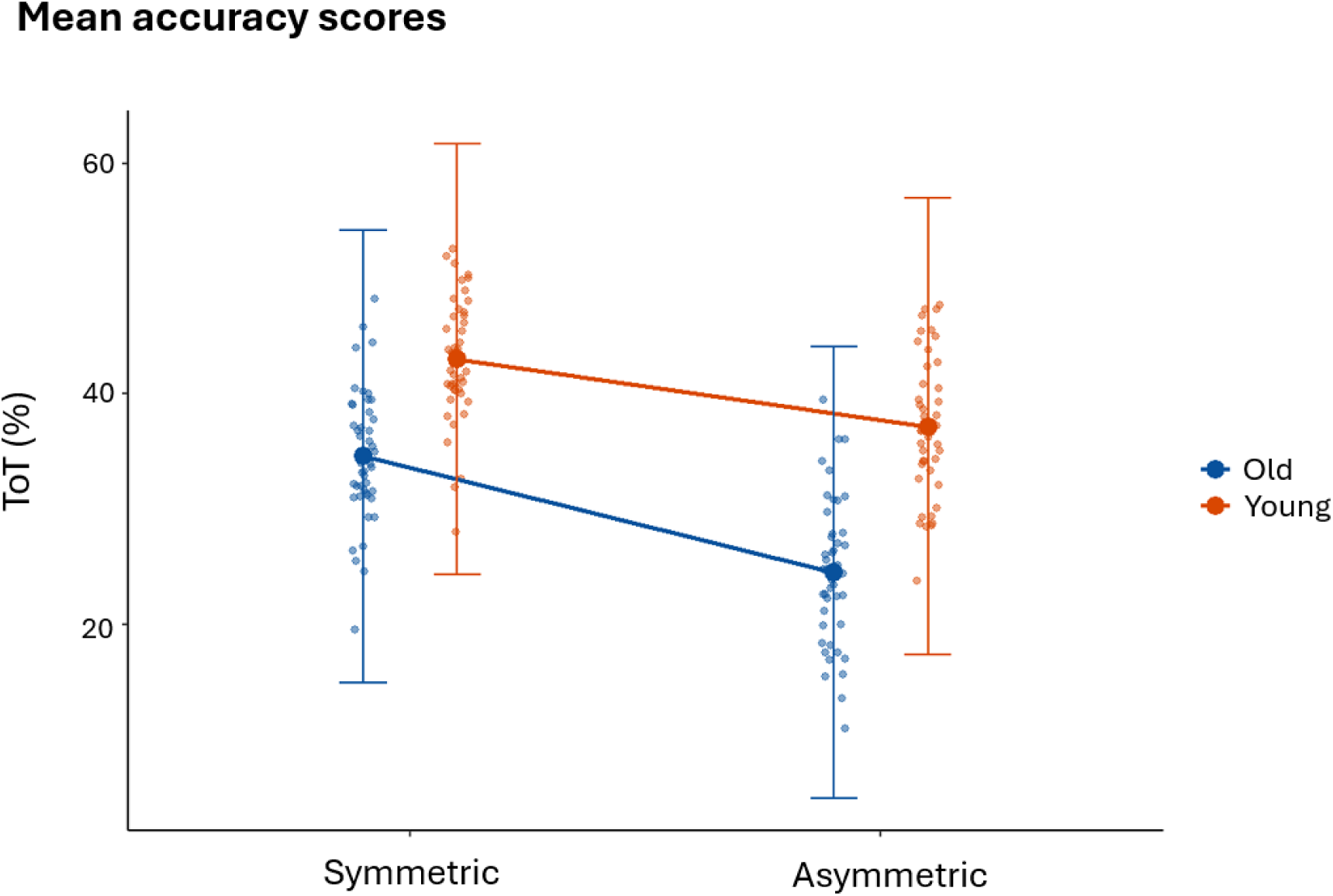
Behavioural results obtained during scanning. Mean accuracy scores, measured as percent time on target (ToT), for young and old participants during the symmetric and asymmetric task conditions. The large dots show the means, while the smaller dots represent individual participants across conditions within their age group.

### 3.2. Neuroimaging results

#### 3.2.1. Main effect of asymmetric and symmetric bimanual movements

The contrast comparing symmetric and asymmetric tasks showed increased bilateral activation during the more complex asymmetric movements in a large bilateral parieto-occipital cluster, as well as in bilateral frontal areas, including the precentral gyrus and large clusters in the prefrontal cortex, as well as the basal ganglia, thalamus, and cerebellum (Figure 3, Table 3). The reverse contrast, focusing on brain areas where symmetric bimanual movements elicited more activation, showed a much more focal activation pattern. Significant clusters were observed in the right precuneus (peak voxel at MNI coordinates [8, −48, 38]; 110 voxels, Z = 3.79) and the left angular gyrus (peak voxel at MNI coordinates [−46, −72, 42]; 112 voxels, Z = 4.91) (Figure 3).

**Figure 3.**
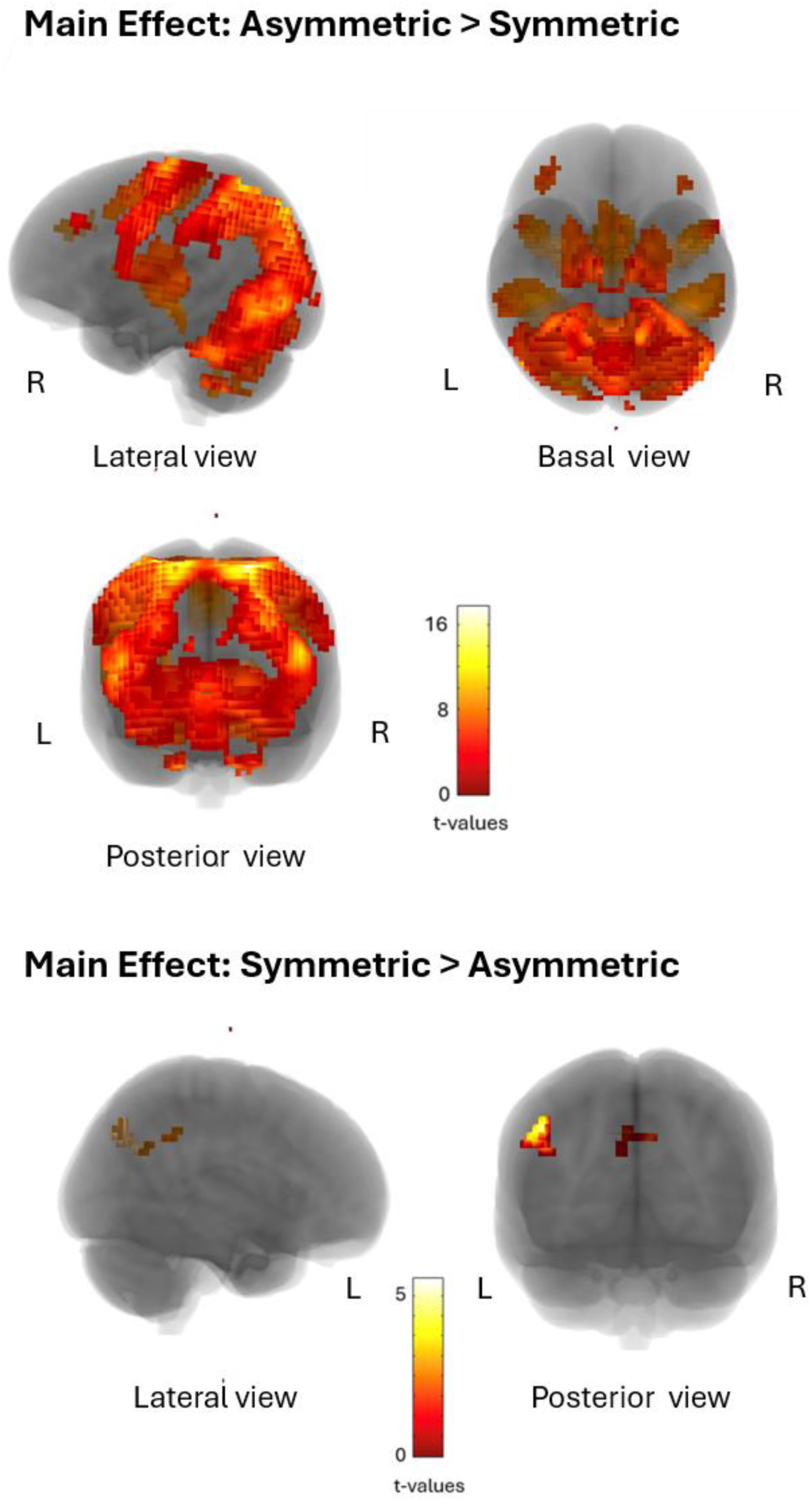
Maximum intensity projections of statistical parametric maps showing an extensive cluster with higher BOLD-signal change for asymmetric bimanual movements compared to symmetric bimanual movements (top) and clusters with higher BOLD-signal change for symmetric bimanual movements compared to asymmetric bimanual movements (bottom) across both age groups. The significance level is set to p<0.05 (FWE) at the cluster-level.

**Table 3:**
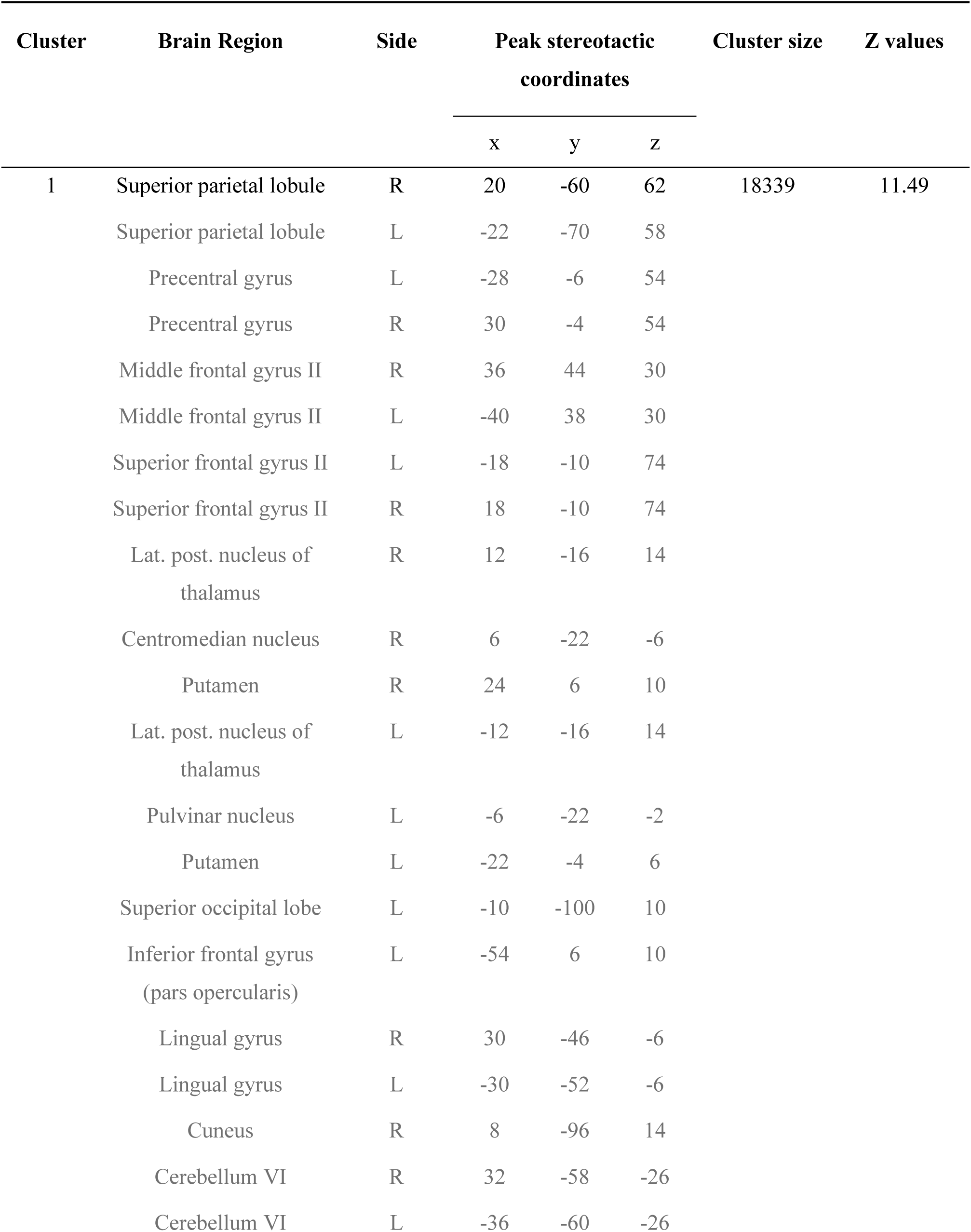
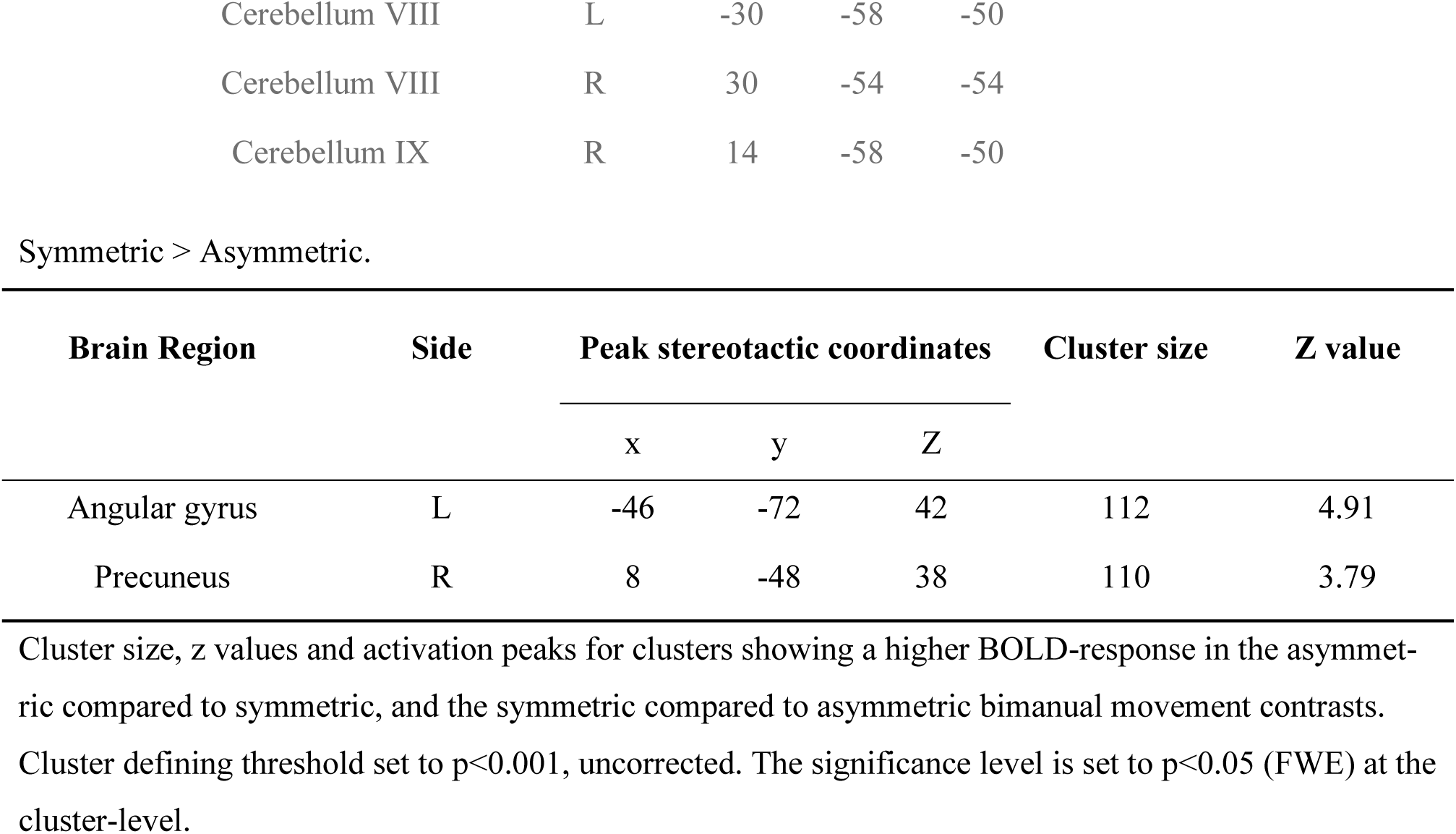
Main effects of task condition for the significant cluster. Asymmetric > Symmetric.

#### 3.2.2. Main Effect of Age Group

The contrast between age groups across tasks revealed only one cluster of higher activation in older adults. The cluster was located in lobules VI and VII of the posterior vermis (peak voxel at MNI coordinates [0, −82, −22]; 465 voxels, Z = 4.47) extending into lobule VI (MNI coordinates [6, −70, −14] and [−4, −72, −14]) and VII in both hemispheres (MNI coordinates [42, −60, −30] and [−31, −58, −29]) (Figure 4). No brain areas were significantly more activated in the younger compared to the older adults.

**Figure 4.**
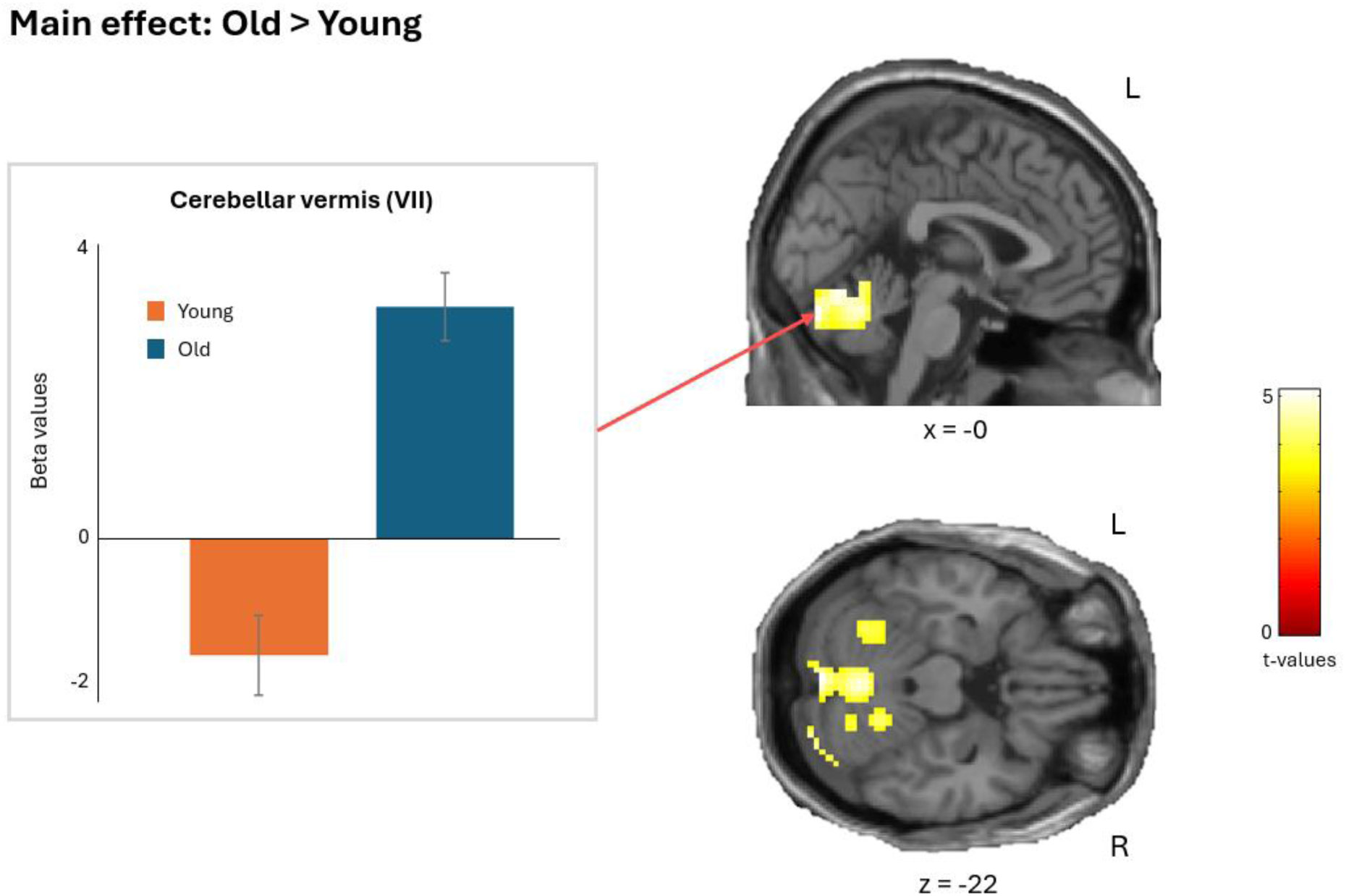
Statistical parametric maps showing a cluster with higher BOLD-signal change for old adults compared to young adults across bimanual task conditions. Cluster defining threshold set to p<0.001, uncorrected. The significance level is set to p<0.05 (FWE) at the cluster-level.

#### 3.2.3. Interaction effect between age and task complexity

The vermis of the anterior cerebellum (lobules IV–V; peak voxel at MNI coordinates [6, −48, −22]; 111 voxels, Z = 4.12), showed a greater increase in activation in the old compared to the young group during the more complex, asymmetric task condition (Figure 5). No other areas showed significantly increased activity during the asymmetric condition in the older adults. The left superior medial frontal gyrus (BA 9/10) (peak voxel at MNI coordinates [−6, 62, 14]; 165 voxels, Z = 3.92) showed the opposite interaction of age group and task type: in this area, younger adults showed increased activation during the more complex asymmetric task condition, while the old group showed decreased activation in this area during the more complex task condition (Figure 5).

**Figure 5.**
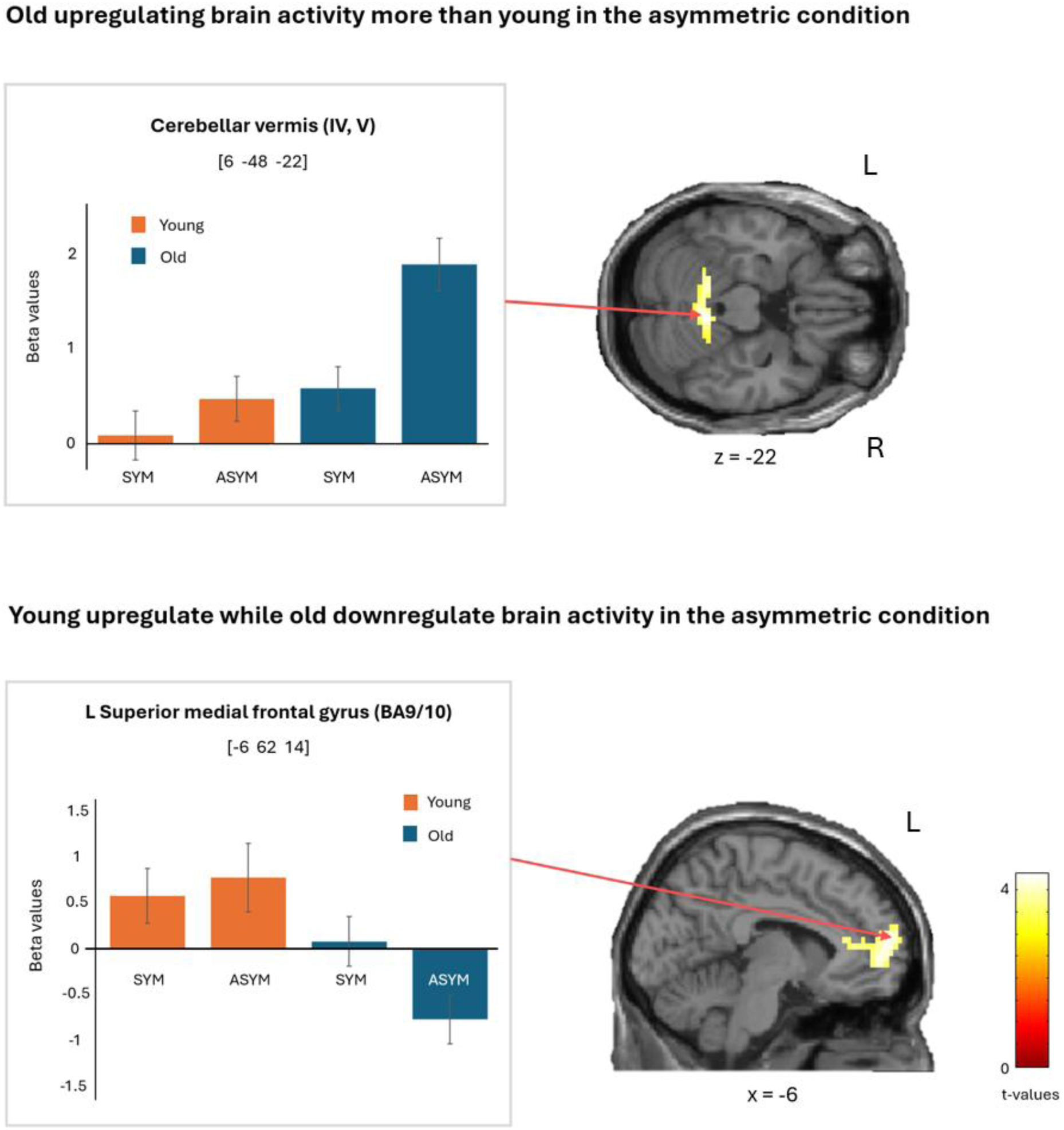
Statistical parametric maps showing a cluster with higher BOLD-signal change in old adults during the more complex asymmetric task condition, and a cluster with higher BOLD-signal change in young adults during the asymmetric condition. Cluster peak beta values are depicted in the bar graphs for each age group and task condition. The significance level was set to p<0.05 (FWE) at the cluster level. Cluster defining threshold set to p<0.001, uncorrected. Error bars show the SE.

#### 3.2.4. Correlation with behavioural performance

The superior medial frontal gyrus (peak voxel at MNI [2, 38, 34]; 165 voxels, Z = 4.85, p = .004, r = −.65 and MNI [14, 50, 2]; 130 voxels, Z = 4.19, p < .01, r = −.58) and right frontal inferior gyrus (peak voxel at MNI [54, 36, 14]; 286 voxels, Z = 4.71, p < .001, r = −.64) correlated negatively with task performance (the greater the brain activation, the poorer the performance) when we combined groups and task conditions (Figure 6). We found no significant correlations between brain activity and task performance when we performed the analysis separately for the groups.

**Figure 6.**
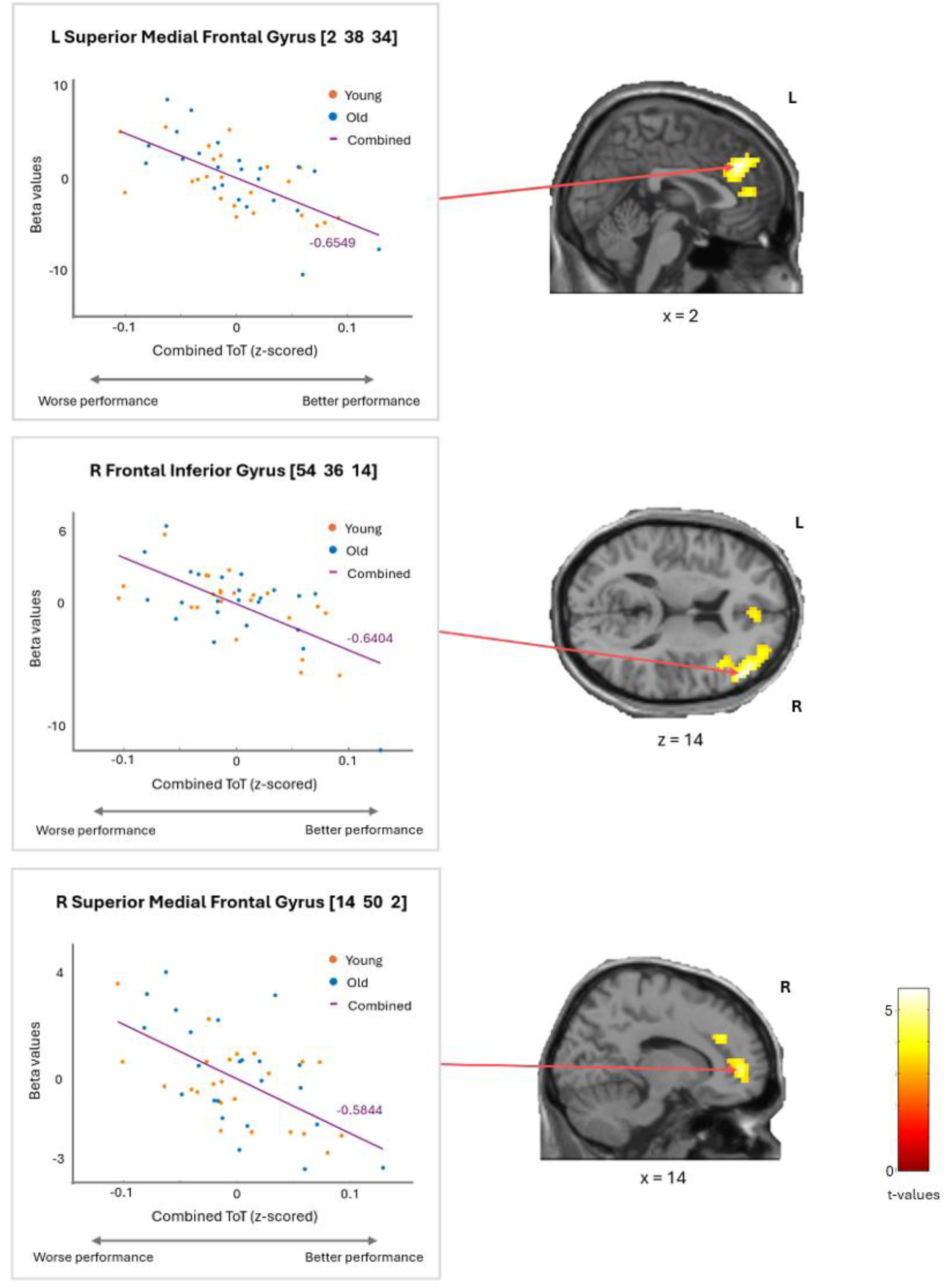
Clusters which correlated negatively with performance for combined age groups and task conditions. Graphs show the correlation between normalized time on target (ToT, z-scored separately within each age group) and beta values for clusters which were significant at the cluster-level p<0.05 (FWE). Cluster defining threshold set to p<0.001, uncorrected.

## 4. Discussion

In this study, we examined whether age-related differences in brain activity during bimanual task performance reflect compensatory over-recruitment of the parieto-frontal regions, as proposed by neurocognitive aging models, or if they reflect a shift towards feedback-mediated motor control processes in older adults. Our results clearly favour the latter. During both simple and complex bimanual coordination, older adults showed impaired task accuracy and increased activation of the posterior vermal lobules (lobules VI-VII), in a cluster overlapping with the classical oculomotor vermis (Stoodley et al., 2012; Voogd et al., 2010). This is consistent with greater reliance on mechanisms involved in visual performance monitoring and online movement correction in older adults, and previous studies have shown that the posterior vermal lobules are involved in error-based adaptation of saccades and movement calibration during visually guided hand movements (Jenkinson & Miall, 2010; Liem et al., 2013; Nitschke et al., 2005; van Broekhoven et al., 2009). During the complex asymmetric coordination task, older adults showed a disproportionately larger decline in performance together with a load-dependent increase in the engagement of the anterior cerebellar vermis (lobules IV–V), in a cluster that is part of the sensorimotor cerebellum (Stoodley et al., 2012). This finding indicates that sensorimotor cerebellum recruitment scaled with task complexity and motor demands, consistent with a greater reliance on online sensorimotor processing when coordinating more challenging movement patterns. We did not observe compensatory over-recruitment of the parieto-frontal regions in older adults in any of the task conditions. Instead, older adults showed decreased activity in the frontal pole (Brodmann area 10) during the asymmetric tasks. Given the established roles of the cerebellum in feedback-based error processing (Ito, 2008; King et al., 2019; Stoodley et al., 2012; Tzvi et al., 2022) and the frontal pole in internal monitoring and goal formation (Burgess et al., 2007; Ramnani & Owen, 2004), this pattern is consistent with a shift toward feedback-driven control in older adults.

### 4.1. Age-related cerebellar recruitment supports increased reliance on feedback-mediated motor control

Across both simple (symmetric) and complex (asymmetric) bimanual tasks, older adults exhibited reduced accuracy and increased activation in the posterior vermal lobules (lobules VI-VII) linking the present findings to a broader literature on visuomotor adaptation, movement calibration, and er-ror-based motor learning (Jenkinson & Miall, 2010; Liem et al., 2013; Nitschke et al., 2005; van Broekhoven et al., 2009). Importantly, this region is also critically involved in visuomotor control and complex bimanual coordination (Boisgontier et al., 2018; Hamano et al., 2025; van Dun et al., 2022), and prior studies have reported age-dependent overactivity in lobules VI-VII during unimanual force-matching tasks (Bower et al., 2025). Structural imaging studies further support the functional relevance of this region for bimanual coordination, as cerebellar grey matter volume in lobule VI is also a strong predictor of bimanual coordination performance across age groups (Boisgontier et al., 2018). Together, these findings indicate that cerebellar lobules VI–VII are central to bimanual motor control and suggest that the higher activation observed in older adults reflects increased engagement of mechanisms involved in sensory prediction error processing, movement calibration, and performance monitoring, consistent with the established role of the oculomotor vermis in saccadic error correction and adaptive motor learning (Jenkinson & Miall, 2010; Tseng et al., 2007; Schlerf et al., 2012). Notably, we did not observe any other significant age-dependent differences in activity between groups, in contrast to prior studies reporting frontal overactivation in regions such as the DLPFC, premotor cortex, or SMA during bimanual tasks (Coxon et al., 2010; Goble et al., 2010; Monteiro et al., 2017), a discrepancy that may reflect the dynamic, non-repetitive nature of our task compared with the more cyclic paradigms used previously. Overall, our findings align with recent meta-analytic evidence highlighting that age-dependent differences are more consistently observed in the cerebellar and occipital-parietal regions than in the frontal cortices during complex bimanual tasks (Zapparoli et al., 2022). These findings indicate that traditional neurocognitive aging models may not generalize to complex bimanual tasks and that age-related patterns of brain activity are more parsimoniously explained by shifts toward feedback-mediated control than by compensatory over-recruitment of the parieto-frontal regions.

Whereas increased activation of posterior vermal lobules VI–VII was observed across task conditions, we also observed that the shift from symmetric to asymmetric coordination affected older adults more than younger adults. Older adults showed a larger drop in task performance and increased activation of more anterior parts of the cerebellar vermis (lobules IV–V) during asymmetric conditions. Lobules IV–V are part of the sensorimotor cerebellum and have been associated with the control of movement timing and execution demands (Manto et al., 2012), which may explain why their activity scales with increasing execution demands during asymmetric coordination, even in the absence of direct performance-activity correlations.

### 4.2. Frontal activity does not support compensatory recruitment

In contrast to the cerebellar findings, we found little evidence that older adults compensated for performance deficits through increased recruitment of frontal control regions. During asymmetric movements, older adults exhibited a decrease in activation of the medial frontal pole (BA10), a region implicated in higher-order cognitive processes including goal maintenance, self-monitoring, and strategy shifting (Gilbert et al., 2006). Reduced recruitment of this region during the most demanding condition may therefore indicate a diminished contribution of higher-order anticipatory control processes, complementing the increased cerebellar engagement observed in regions associated with feedback-related motor processing.

Similarly, analyses of performance-related activity revealed that better task performance was associated with decreased activation in the superior medial frontal gyrus and right inferior frontal gyrus across both age groups. These areas are commonly implicated in cognitive control and response selection/inhibition (Aron et al., 2014; Levy & Wagner, 2011). Importantly, the observed relationships were not age-specific but were present across younger and older participants. Rather than supporting compensatory recruitment, these findings suggest that successful task execution was associated with reduced engagement of frontal cognitive control mechanisms. Together, the absence of frontal overactivation in older adults, the reduced recruitment of BA10 during the most demanding condition, and the negative association between frontal activity and task performance, argue against neurocognitive aging models that predict compensatory frontal recruitment. Instead, the findings support the notion that age-related differences in task performance are more closely linked to altered cerebellar and sensorimotor processing than to increased reliance on frontal executive resources.

### 4.3. Task-dependent differences in activity

Generally, we found that both age groups had increased activity across a wide range of brain regions during the asymmetric movements. These regions included the sensorimotor areas, supplementary motor area, inferior frontal gyrus, basal ganglia, and the posterior cerebellum, a pattern similar to that reported in other studies when shifting from simple to complex bimanual tasks (Coxen et al., 2010; Goble et al., 2010; Karabanov et al., 2023; Van Ruitenbeek et al., 2023). This pattern is consistent with increased demands during asymmetric bimanual coordination requiring the decoupling of default symmetric motor patterns and the generation of distinct internal timing models for each limb, increasing reliance on cerebellar forward modelling and parietal sensorimotor integration.

Greater activity in the symmetric compared to the asymmetric condition was observed in the left angular gyrus and right precuneus. Both regions are central to higher-order visuospatial integration and the representation of global spatial relationships (Mahayana et al., 2014; Wagner & Rusconi, 2023). Symmetric bimanual movements allow the two hands to be coordinated as a single coherent spatial pattern, placing greater emphasis on integrated visuospatial representations rather than limb-specific predictive control. Increased activation in the precuneus and angular gyrus during the symmetric condition may therefore reflect reliance on global spatial integration and automatized coordination, which are reduced when movements become asymmetric and require independent timing and prediction for each limb.

### 4.4. Limitations and future directions

A known limitation of fMRI in aging research is that the blood-oxygen-level-dependent (BOLD) signal reflects both vascular and neuronal contributions, and these vascular factors change with age. Age-related alterations in cerebrovascular structure and neurovascular coupling can influence BOLD amplitude and timing independently of neuronal activity, potentially confounding group comparisons of functional activation patterns (Stiernman et al., 2023; Tsvetanov et al., 2021).

White matter hyperintensities (WMH), which are more prevalent in older adults and reflect small-vessel pathology, have been linked to reduced cerebral blood flow and are therefore a potential indirect proxy for vascular health and regional vascular compromise in the older group (Marstrand et al., 2002; Wardlaw el al., 2013). In the present study, we have taken several precautions to mitigate vascular confounds. We corrected for physiological variables such as heart rate and respiration that influence BOLD signal fluctuations and controlled for WMH burden as an indirect index of regional vascular compromise. This combination of physiological denoising and structural adjustment should reduce non-neuronal contributions to BOLD signal differences between age groups, strengthening the inference that observed activation patterns reflect underlying neural processing rather than purely vascular differences.

## 5. Conclusion

We found that during performance of a bimanual task, older adults, compared to young adults, showed increased activation of posterior cerebellar regions across task conditions and stronger recruitment of the anterior cerebellar vermis as task complexity increased. This pattern of brain activation suggests that age-related differences in the neural processes underlying task performance reflect a shift towards feedback-mediated motor control rather than an increased reliance on frontal executive resources. In contrast, we found little evidence for compensatory frontal recruitment.

Older adults showed reduced activation of the frontal pole during the most demanding condition, and better task performance across task conditions was associated with lower rather than higher frontal activation across both age groups. Together, these findings suggest that age-related differences in brain activity, during complex bimanual coordination, are more closely linked to altered cerebellar and sensorimotor processing than to compensatory recruitment of frontal executive resources.

## Acknowledgements

Anke Karabanov has received funding as principal investigator for the project “Reconfigurations in Large-Scale Brain Networks - ReScale” from Danmarks Frie Forskningsfond (grant nr. 0169-00027B). Hartwig R. Siebner has received funding as principal investigator for the project “Precision Brain-Circuit Therapy - Precision-BCT” from Innovation Funds Denmark (grant nr. 9068-00025B) and the project “ADAptive and Precise Targeting of cortex-basal ganglia circuits in Par-kinsońs Disease - ADAPT-PD” from Lundbeckfonden (collaborative project grant, grant nr. R336-2020-1035).

The authors are thankful to Keenie Ayla Andersen and Ana Zvornik for helping with data acquisition. Authors’ contributions: conceptualization: AK, MN, KHM, JL, HRS; funding acquisition: AK; data collection: MN; data analysis: ASW, MN, KHM; supervision: MN, AK; Visualization: ASW, MN; Writing – original draft: ASW, AK; Writing – review and editing: ASW, MN, KHM, JL, HRS and AK.

