## Supplementary 2 for "Age-Related Differences in Bimanual Coordination Are Associated with Increased Cerebellar Activity and Reduced Frontal Recruitment"

**Supplementary 2:** Neuroimaging results for the two interaction clusters showing the % change between symmetric and asymmetric task conditions.

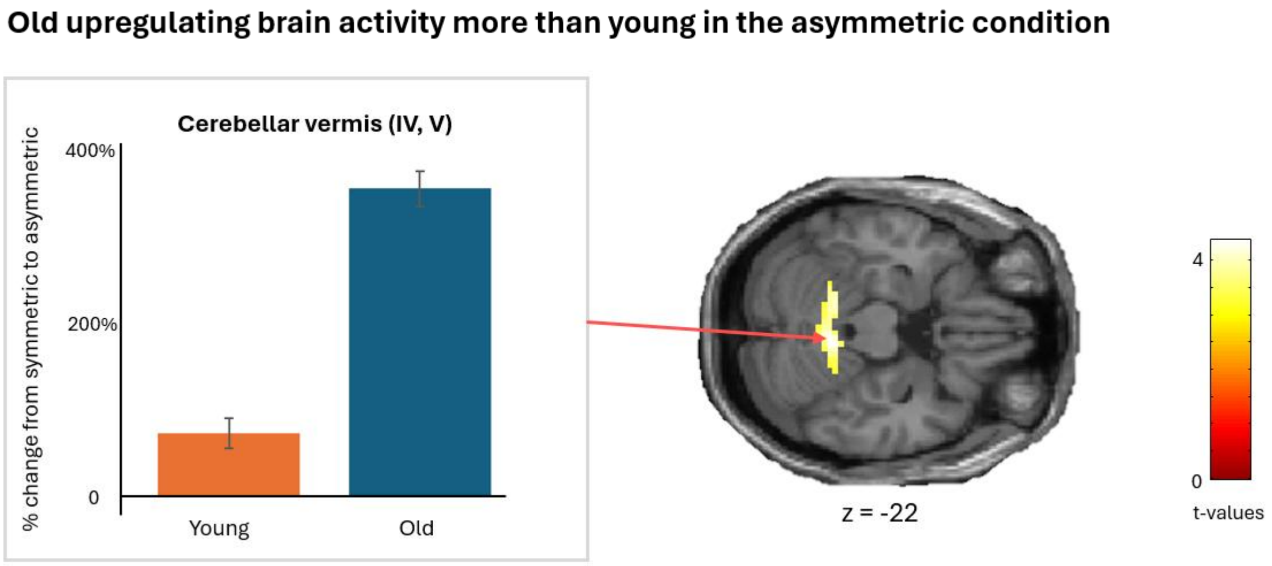


**Figure 7.** Statistical parametric map showing a cluster with higher BOLD-signal change in old adults during the more complex asymmetric task condition. Beta values are depicted in the bar graphs for each age group and task condition. The significance level is set to $p < 0.05$ (FWE) at the cluster-level. Error bars show the SE.

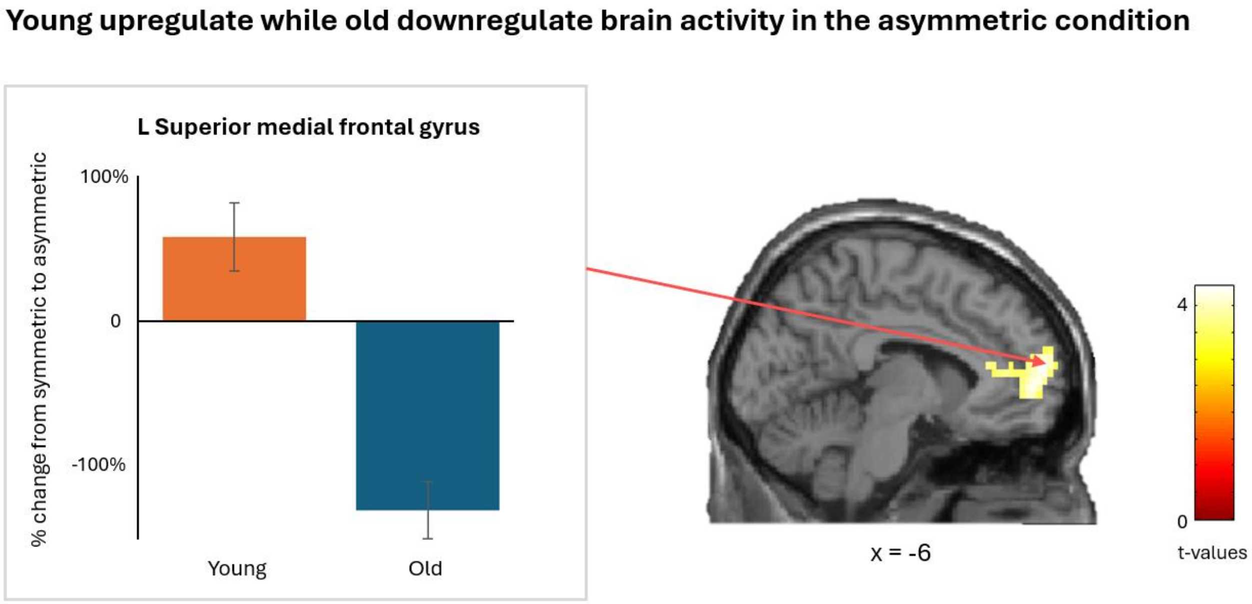


**Figure 8.** Statistical parametric map showing a cluster with higher BOLD-signal change in young adults during the more complex asymmetric task condition. Beta values are depicted in the bar graphs, showing the difference between task conditions for each age group. The significance level is set to $p < 0.05$ (FWE) at the cluster-level. Error bars show the SE.
